# Changes in rearing conditions rapidly modify gut microbiota structure in *Tenebrio molitor* larvae

**DOI:** 10.1101/423178

**Authors:** Marine Cambon, Jean-Claude Ogier, Anne Lanois, Jean-Baptiste Ferdy, Sophie Gaudriault

## Abstract

The gut microbiota of multicellular organisms has been shown to play a key role in their host biology. In mammals, it has an invariant component, responsible for establishing a mutualistic relationship with the host. It also contains a dynamic fraction which facilitates adaptation in response to changes in the environment. These features have been well described in mammals, but little is known about microbiota stability or plasticity in insects. We assessed changes in microbiota composition and structure in a reared insect after a change in rearing conditions. We reared *Tenebrio molitor* (*Coleoptera, Tenebrioninae*) larvae for five days in soil samples from two river banks and analyzed their gut microbial communities by a metabarcoding technique, using the V3-V4 region of the 16S rRNA gene and the housekeeping gene *gyrB*. We found that soil-reared insects had a significantly more diverse microbiota than the control insects and that insects reared in soil from different sites had significantly different microbiota. We confirmed this trend by absolute quantification of the two mains fluctuating taxonomic groups: the *Enterobacteriaceae* family and the *Pseudomonas* genus, dominant in the soil-reared insects and in the control insects, respectively. Our results suggest the existence of a resident microbiota in *T. molitor* gut, but indicate that rearing changes can induce rapid and profound changes in the relative abundance of some of the members of this resident microbiota.

## Background

Microorganisms have repeatedly been shown to play a key role in plant and animal biology (Bordenstein and Theis 2015). If we are to understand the biology of a pluricellular organism, we must consider its microbiota, the cohort of microorganisms associated with the host. In animals, the gut microbiota is a key component, with major effects on host physiology. For example, the mammalian gut microbiota has been the object of many studies on digestive functions with health implications (Belizário and Napolitano 2015).

The composition of the mammalian gut microbiota displays both plasticity and invariant features. The core microbiota, which consists of the microorganisms common to the majority of individuals within a population, is generally defined as the most prevalent of the microbial species detected (Shetty et al. 2017). This common fraction of the microbiota plays a fundamental role in supporting the mutualistic symbiotic relationship with the host (Candela et al. 2012). For example, changes in the human core microbiota are associated with physiological perturbations, such as obesity and Crohn’s disease (Turnbaugh et al. 2009; Hedin et al. 2015). However, another key feature of the mammalian gut microbiota is its plasticity, i.e. its ability to change in composition and structure. In humans, dietary changes induce a remarkable degree of variation in gut microbiota in terms of both phylogenetic and functional composition (Candela et al. 2012). These changes depend on various factors including host age, sex, genetic make-up, immune and health status (Shetty et al. 2017), but also exposure to environmental bacteria, geographic origin and climate (Candela et al. 2012). It has been suggested that this plasticity of the human gut microbiota facilitates rapid responses to environmental change, resulting in rapid ecological adaptation (Alberdi et al. 2016).

Most studies on the gut microbiota concern mammals. However, the use of mammals, and more generally of vertebrates, in experimental approaches raises numerous practical, financial and ethical issues. Large-scale experiments require model organisms that are easy to manipulate and can be obtained in large numbers. Insects are interesting experimental models in this respect. Although their guts contain fewer microbial species than those of mammals (Engel and Moran 2013), insects also rely on their gut microbiota for diverse functions, including development, nutrition, the modulation of immune responses, gut homeostasis, protection from pathogens and toxins (Engel and Moran 2013; Shi et al. 2013; Broderick et al. 2014; Erkosar and Leulier 2014; Caccia et al. 2016; Welte et al. 2016; Shao et al. 2017; Raymann and Moran 2018). The gut microbiota of non-social insects is principally acquired from the environment through feeding (Engel and Moran 2013). Its composition depends on environmental conditions and diet in both laboratory and wild individuals (Chandler et al. 2011; Montagna et al. 2015; Staudacher et al. 2016). For example, it has been shown for some coleopteran species that microbiota changes with geographical location, environmental condition, and life stage (Huang and Zhang 2013; Montagna et al. 2014).

One potential limit of these previous studies is that they used either insects from the wild, which cannot be controlled for many of their characteristics, or lab-reared insects, which are controlled but have a poorly diversified microbiota. Here we used laboratory-reared *T. molitor* larvae and mimicked a soil environment by rearing the larvae for five days in different soil samples. We assessed the changes in gut microbiota composition after acclimatization to soil samples and demonstrated a large shift in gut microbial structure. We showed in addition that different soil samples induced different modifications in insect microbiota, and that the observed plasticity was probably dependent on changes in the abundance of some of the resident OTUs.

## Methods

### Soil samples

We sampled soil from riverside land around Montpellier in the South of France (Figure 1A): on the banks of the Hérault river near Causse-De-La-Selle (N43°49.884′ E003°41.222′; CDS sample) and on those of the Lez river near Montferrier-sur-Lez (N43°40.801’ E003°51.835’; MTF sample). Both soils had a sand-silt-clay composition typical of riversides on chalky substrata. The sand:silt ratio was higher for MFT than for CDS. We collected three soil subsamples from each plot. These subsamples were taken at a depth of 20 cm and were separated by a distance of 10 m. They were named CDS1, CDS2, CDS3 and MTF1, MTF2, MTF3 (Figure 1B). The use of these six soil subsamples made it possible to compare the variability in microbiota composition both between and within plots. Each soil subsample was split into four portions, each of which was placed in a 1 L plastic box (Figure 1C), in which it was mixed with heat-sterilized (20 min at 121 °C) wheat bran (1:3 (v/v) ratio, as previously described (Jung et al. 2014).

**Figure 1:**
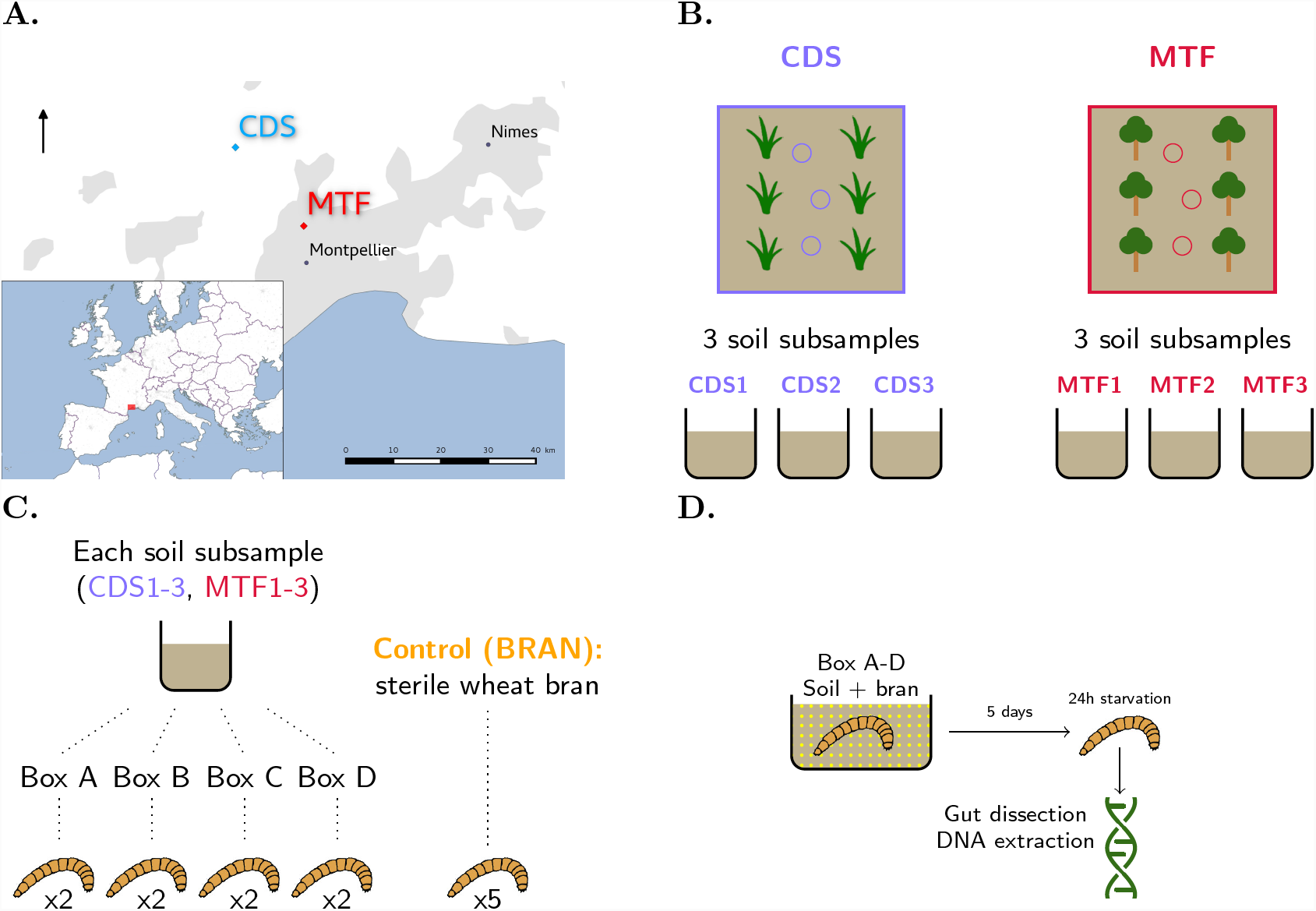
Experimental design. **A.** Location of the two sampling sites. CDS: Causse-De-La-Selle (N43°49.884’ E003°41.222’; CDS sample); MTF: Montferrier-sur-Lez (N43°40.801’ E003°51.835’; MTF sample). **B.** At each sampling site, we obtained three soil subsamples at positions 10 m apart. **C.** Distribution of insects in soil subsamples. Each soil subsample was split into four portions, each of which was placed in a plastic box, in which it was mixed with sterilized wheat bran. Eight insects per soil subsample (two insects/box) were analyzed. Five insects placed in a box containing sterile wheat bran only were used as a control. **D.** Insects were placed, for five days, at 15 °C, in plastic boxes containing the soil subsamples mixed with sterile wheat bran. They were then starved by incubation for 24 hours in Petri dishes. The insects were then killed, their guts were dissected, and total DNA was extracted from each gut.

### Insects

Larvae were provided by Micronutris (St-Orens, France) and fed with heat-sterilized bran before the experiment. As it was not possible to determine their precise developmental stage, but we used only larvae weighing between 20 and 50 mg, which should correspond to 13th or 14th instar individuals (Huang et al. 2011).

### Rearing of *T. molitor* larvae in soil samples

We maintained laboratory-reared *T. molitor* larvae for five days in sterilized wheat bran mixed with soil samples. During this period, the larvae were incubated at 15 °C in the same humidity conditions. They were then starved for 24 hours (Figure 1D) to exclude individuals that were infected with pathogens (which would have died within this 24 hours period) and to limit the risk that the DNA we extract comes from the larval alimentary bolus. This starvation period potentially induces a stress on insect larvae, which might in turn impact their microbiota. We imposed it on all insects, so that the potential bias it creates is identical in all treatments.

Control insects were reared in the same conditions than other insects except that they were incubated in sterile wheat bran, with no soil mixed. Control insects microbiota should thus be close to what it was for all insects before the experiment.

### DNA extraction

We extracted DNA from two randomly sampled insects per box (which makes a total of 24 insects per site) and 5 control insects. However, we failed to amplify 16S rRNA during PCR step for 2 samples, ending with 24 samples for CDS, 22 samples for MTF and 5 controls. Insect larvae were sterilized in 70% ethanol, rinsed in water and then killed. The guts of the larvae were dissected in sterile Ringer solution (Jung et al. 2014). Dissection tools were sterilized with 70% ethanol between insects. Dissected guts were placed in an Eppendorf tube with 100 μL of lysis solution and 1 μL lyzozyme (Quick Extract, Bacterial DNA extraction TEBU-BIO) and ground with 3 mm steel beads for 30 seconds at 20 Hz with a TissueLyzer (Qiagen). The resulting homogenates were incubated at room temperature for two days, then frozen in liquid nitrogen and heated at 95 °C to ensure that all the cells were lysed. DNA was prepared by the phenol-chloroform-alcohol and chloroform extraction method. The DNA was resuspended in sterile water and quantified with a NanoDrop spectrometer (Thermo Fisher Scientific). We performed extraction blanck controls using DNA-free water.

### 16S and *gyrB* DNA amplification

We targeted the V3-V4 region of the 16S rRNA gene, which is classically used for bacterial identification in microbial ecology studies, as clean and complete reference databases are available for this region. We also used the bacterial housekeeping gene *gyrB*, to support the data for the 16S rRNA (Barret et al. 2015). The V3-V4 region of the 16S rRNA gene was amplified with the PCR1F_460 (5’-ACGGRAGGCAGCAG-3’) / PCR1R_460 (5’-TACCAGGGTATCTAATCCT-3’) primers (modified versions of the primers used in a previous study Klindworth et al. (2012)). Amplification was performed with the MTP Taq polymerase (Sigma, ref 172-5330), according to the manufacturer’s protocol, with 1 μL of 1/10 diluted DNA extract for each sample. The PCR protocol used for these primers was 60 s at 94 °C, followed by 30 cycles of 60 s at 94 °C, 60 s at 65 °C, 60 s at 72 °C, and then 10 min at 72 °C. The *gyrB* gene was amplified with primers described elsewhere: gyrB_aF64 5’-MGNCCNGSNATGTAYATHGG-3’ and gyrB_aR353 5’- ACNCCRTGNARDCCDCCNGA-3’ (Barret et al. 2015). Amplification was performed with the iProof High-Fidelity Taq polymerase (Bio-Rad, ref. 172-5301), according to the manufacturer’s protocol, with 1 μL of 1/10 diluted DNA extract for each sample. The PCR protocol used for these primers was 30 s at 98 °C, followed by 40 cycles of 10 s at 98 °C, 30 s at 60 °C, 30 s at 72 °C, and then 10 min at 72 °C. For each PCR, we performed negative and positive controls with water and bacterial DNA extracted from a pure culture of *Xenorhabdus nematophila* (*Enterobacteriaceae*), respectively, and checked PCR amplicons by electrophoresis in a 1% agarose gel. We performed technical replicates for the PCR and sequencing steps and obtained identical microbiota patterns (see Additional File 2, for example). Amplicon libraries were sequenced by the GeT-Plage genomics platform at Genotoul (Toulouse, France) with Illumina MiSeq technology and a 2×250 bp kit. Raw sequence data of both 16S rRNA and *gyrB* are available from http://www.ebi.ac.uk/ena/data/view/PRJEB21797.

### Metabarcoding data treatment

Sequence data for both markers were analyzed with OBITools (Boyer et al. 2015). Raw paired-end reads were aligned and merged, taking into account the phred quality score of each base to compute an alignment score. Reads with a low alignment score (>50), containing unknown bases or with an unexpected size (outside 400 bp and 470 bp, and 230 bp and 260 bp after primer trimming for the 16S rRNA gene and *gyrB* respectively) were removed from the dataset. After primer trimming, singletons (i.e. sequences only found once in the dataset) were removed (Auer et al. 2017). Sequences were then clustered into OTUs with the Sumaclust algorithm (Mercier et al. 2013), using a 97% similarity threshold (OBITools workflows and the raw count table are available in Additional Files 3 and 4). We then removed from the datasets all clusters containing less than 0.005% of the total number of reads (Bokulich et al. 2013). The remaining OTUs were assigned to a taxonomic group with RDPclassifier (Wang et al. 2007) and the RDPII reference database for the 16S rRNA marker and with seq_classifier.py from the mothur pipeline (Schloss et al. 2009) and the reference database from Barret et al. (2015) for *gyrB* gene (OTU assignments are available in Additional File 5).

### Quantitative PCR analysis

To check for changes in OTU abundances, we performed quantitative PCR (qPCR) on two randomly picked insects per soil subsample among those used in the metabarcoding analysis. The sampling probability for each sample was adjusted for the total number of 16S rRNA reads for the sample. The five DNA samples corresponding to control insects were all analyzed.

All qPCRs were performed in triplicate, with 3 μL of reaction mixture, on a LightCycler480 machine (Roche Diagnostics), after the plate had been filled by an Echo 525 liquid handler (Labcyte). The reagent concentrations were identical in all SYBR Green I assay reactions: 1×Light-Cycler 480 SYBR Green I Master Mix (Roche Diagnostics), 500 nM each of the forward and reverse primers specific for genus *Pseudomonas* (here named *Pse* -16S, Bergmark et al. (2012)), the *Enterobacteriaceae* family (here named Entero-*rplP,* Takahashi et al. (2017)) or the *Eubacteria* kingdom (here named *uni16S,* Vandeputte et al. (2017)) (see sequences in Additional File 6) and DNA matrix. The DNA used was either genomic DNA (0.5 ng/μL) from the various reference strains, to check primer specificity (*Escherichia coli, Serratia marcesens, Klebsiella pneumoniae, Salmonella typhimurium, Enterobacter cloacae, Pseudomonas protegens, Stenotrophomonas, Acinetobacter, Enterococcus*) or a 1/100 dilution of insect gut DNA for metabarcoding. The qPCR conditions were 10 minutes at 95 °C, followed by 45 cycles of 5 s at 95 °C, 10 s at 62 °C and 15 s at 72 °C, with a final dissociation curve segment. Cycle threshold (Ct) values were determined with Light-Cycler 480 software. After the validation of primer specificity (13 < Ct < 37 for positive controls, Ct > 40 for negative controls), absolute quantifications were calculated by the standard curve method. Serial dilutions of standard samples consisting of genomic DNA from *E. coli* ATCC25922 for the *rplP* gene and the rRNA16S gene (*uni16S* primers) and genomic DNA from *Pseudomonas aeruginosa* CIP76.110 (=ATCC27853) for the 16S rRNA gene (*Pse* -16S primers) were prepared and used for calibration. The gene copy number of the target gene (*GC N*_*target*_ [copies]) in standard samples was estimated using the total amount of genomic DNA in the calibration samples *M*_*DNA*_ [g], the size of the bacterial chromosome *L*_*DNA*_ [bp], the number of targets per bacterial chromosome *n*_*target*_ [copies], Avogadro’s constant *N*_*A*_ (6.022 × 10^23^ bp mol^-1^) and the mean weight of a double-stranded base pair *M*_*b p*_ (660 g mol^-1^bp^-1^) as follows:

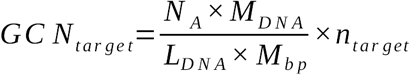

Using the parameters of the curves linking *GCN*_*target*_ and *C t* in standard samples, we then estimated the GCN of target genes in our gut samples. This estimation was possible because PCR efficiency (PE) was very close to that for standard samples (Additional File 6).

### Community analysis

All analyses were performed with R version 3.3.3 (R Core Team 2015) (see Additional File 7 and 8 for the overall workflow). We did not rarefy data (McMurdie and Holmes 2014), but we used Chao1 index which is the estimated OTU richness of each sample, taking into account the possible lack of detection of some rare OTUs. Chao1 index is thus the observed OTU richness per insect plus an estimation of the unseen OTUs per insect. The Shannon index is based on relative abundance data, to represent the effective OTU richness of the sample based on the predominant OTUs. We estimated the Chao1 and Shannon alpha diversity indices with the vegan package of R (Oksanen et al. 2017). We also calculated Pielou’s eveness which is the Shannon diversity divided by the natural logarithm of the OTU richness of the sample, and reflects how similar the relative abundances of OTUs in a sample are.

We calculated the beta diversity distance matrix from the Jaccard and Bray-Curtis distances for presence/absence and relative abundance data, respectively, using the vegan package. We also computed Unifrac and wUnifrac distances for presence/absence and relative abundance data, respectively (Lozupone and Knight 2005), with the Phyloseq package (McMurdie and Holmes 2013). Unifrac and wUnifrac distances include phylogenetic distances between pairs of OTUs. A phylogenetic tree of the OTU sequences was, therefore, required. We generated this tree by aligning OTU sequences with Seaview software and the muscle method. The phylogenetic tree was built with RA×ML and the GTRCAT substitution model for nucleotide sequences (Stamatakis 2014) (Additional File 9). Differences in the gut bacterial community between soil-reared insects and control insects were evaluated based on the beta diversity distance matrix, in PERMANOVA tests implemented in the vegan package (Oksanen et al. 2017), with treatment as the explanatory variable. We investigated differences between the gut bacterial communities of soil-reared insects, by performing PERMANOVA tests on distance matrices with two explanatory variables: soil sample (i.e. CDS or MTF) and soil subsample (i.e. CDS1-3, MTF1-3). Beta-diversity distances were represented using a PcoA analysis from the vegan package (Oksanen et al. 2017).

## Results

### Incubation of *T. molitor* larvae with soil increases the richness and diversity of their gut microbiota

After cleaning, the total dataset of the metabarcoding experiment contained 792,395 sequences clustered into 106 bacterial OTUs. Rarefaction curves showed that most of the samples had reached the saturation plateau (Figure 2A). We used the Chao1 index, which assesses the extrapolated richness of OTUs, including an estimation for undetected rare OTUs, to compare alpha diversity between soil-reared and control insects. The mean Chao1 index of the microbiota of soil-reared insects (MTF and CDS) was a 48 *±* 13 OTUs whereas that of control insects (BRAN) was 25 *±* 9 OTUs (Figure 2B). The OTU richness of the gut microbiota therefore increased significantly after the incubation of the insects with soil samples (Chao1 index, soil vs. control: Wilcoxon rank sum test, W=221, p-value = 1e-3). A similar conclusion was drawn for analyses based on the Shannon index, which reflects relative OTU abundance within samples (Figure 2B, soil vs. control: Wilcoxon rank sum test, W=216, p-value = 1e-3). Moreover, control insects harbored bacterial communities dominated by a very small number of dominant OTUs (low Shannon index *≃* 0.2 and low Pielou’s eveness *≃* 0.02). OTU assignment identified these dominant OTUs as belonging to the *Pseudomonadaceae* family (Figure 2C). By contrast, soil-reared insects harbored bacterial communities with more balanced relative OTU abundances (Shannon index *≃* 1.7). The gut microbiota of these insects was dominated by *Enterobacteriaceae*, together with *Pseudomonadaceae* and other less frequent families, such as *Moraxellaceae* and *Aeromonadaceae* (Figure 2C). This was confirmed by the analysis of Pielou’s eveness index which was significantly lower in control insects than in soil-reared insects (Wilcoxon rank sum test, W=0, p-value = 7.6e-7). Thus, five days in soil significantly increased the richness of the microbiota in the gut of *T. molitor* larvae, and modified the balance of OTUs present.

**Figure 2:**
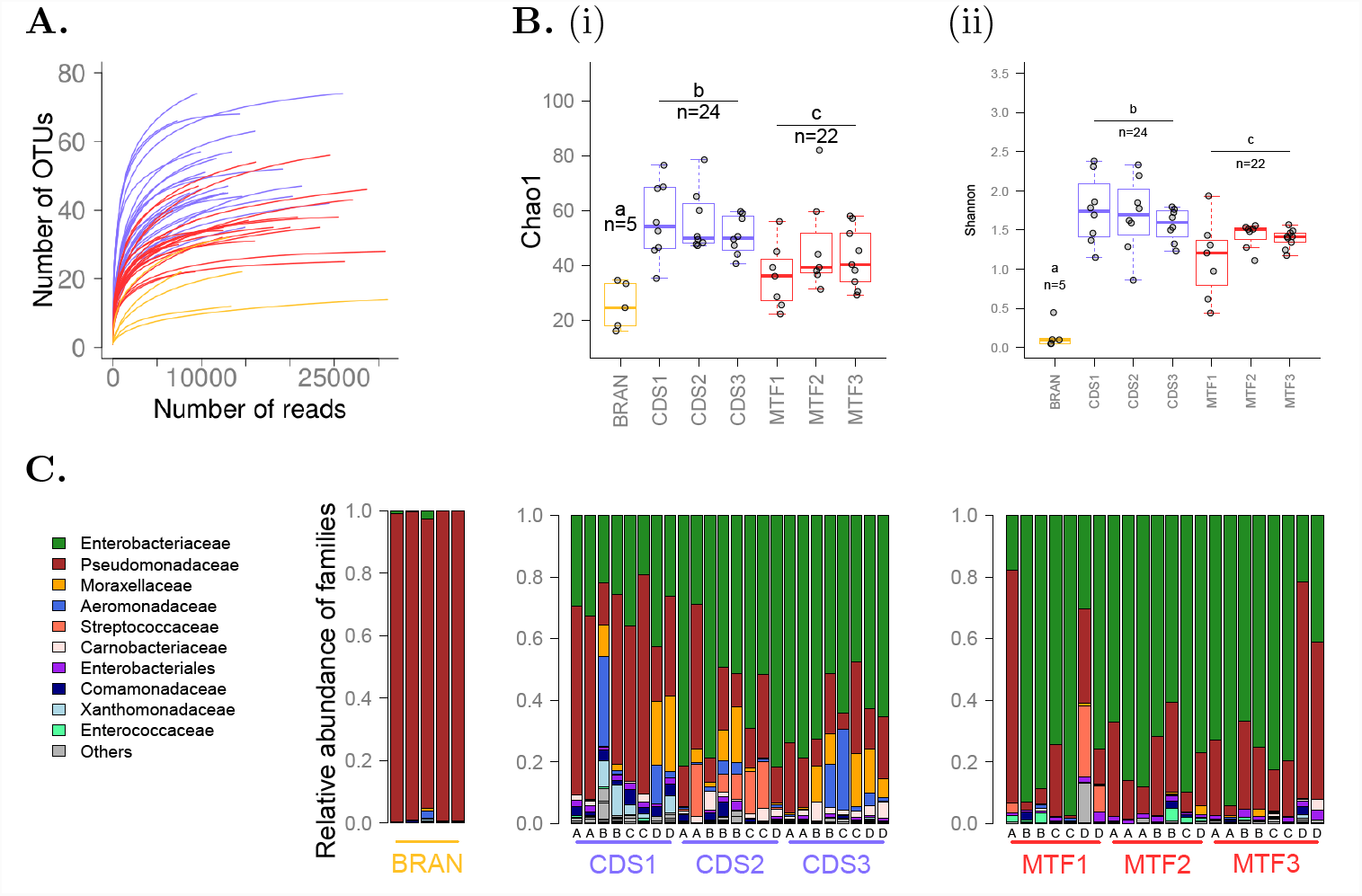
Alpha diversity of the gut microbiota. **A.** Rarefaction curves. Each curve represents one insect. Control insects, insects reared in CDS soil samples and insects reared in MTF soil samples are shown in yellow, blue and red, respectively. **B.** Alpha diversity indices for the insect gut microbiota. CDS1-3 and MTF1-3 are the subsamples from the sampling sites (three for CDS and three for MTF). BRAN is the control treatment: insects reared on sterile wheat bran. (i) Chao1 extrapolated richness. Pairwise Wilcoxon rank-sum test, CDS-MTF: p-value = 2e-3, BRAN-CDS: p-value = 2e-3, BRAN-MTF: p-value = 0.01 (ii) Shannon diversity index. Pairwise Wilcoxon rank-sum test, CDS-MTF: p-value = 1e-3, BRAN-CDS: p-value = 5e-05, BRAN-MTF: p-value = 8e-05 **C.** Taxonomic assignment of OTUs to family level. Each bar represents an insect. Each subsample (i.e. CDS1-3 and MTF1-3) was divided into four portions, each of which was placed in a separate plastic box before the experiment. For each subsample, insects sharing the same letter (A, B, C or D) were taken from the same plastic box. The 10 families with the largest relative abundances are shown in different colors, and the others are grouped together in the “Others” category.

We also investigated the effect of soil treatments according to soil origin, by comparing the alpha diversity of CDS and MFT samples. The Chao1 and Shannon indices were significantly lower in MTF than in CDS samples (Figure 2B; Chao1 index: Kruskal-Wallis test, *χ*^2^=12.93, p-value = 3e-4; Shannon index: Kruskal-Wallis test, *χ*^2^=9.6136, p-value = 1e-3). The CDS and MTF soils had therefore different impacts on both richness and bacterial balance.

### Soil treatment induces a change in microbiota composition that is variable between soil sampling sites

We investigated the effect of soil treatment on insect microbiota, by calculating the beta-diversity between insect gut microbiota with various distance indices (Figure 3). We first calculated a distance based on pairwise Jaccard and Bray-Curtis distances. These two indices are complementary, because Jaccard distance depends purely on the presence/absence of OTUs, whereas Bray-Curtis distance also takes into account the number of reads for each OTU as a proxy for their relative abundance. We performed PCoA analysis on distance matrices (Figure 3A) where control insects tended to cluster together. PERMANOVA analysis confirmed that community composition differed between soil-reared insects and control insects (13 to 19% of the variance explained by soil treatment, Table 1A).

**Table 1:** PERMANOVA analysis of the community composition of the insect microbiota based on different diversity indices, with the percentage of the variance explained by each variable and the p-value in brackets

| Variable | Jacc | BC | Uni | wUni |
| --- | --- | --- | --- | --- |
| **A.** | *All insects* |  |  |  |
| Treatment | 0.13 (1e-3) | 0.19 (2e-3) | 0.03 (0.07) | 0.18 (1e-3) |
| **B.** | *Soil-reared insects* |  |  |  |
| Site | 0.14 (1e-3) | 0.08 (2e-3) | 0.09 (1e-3) | 0.07 (6e-3) |
| Subsample | 0.17 (1e-3) | 0.18 (1e-3) | 0.14 (3e-3) | 0.20 (1e-3) |

Jaccard distances (Jacc), Bray-Curtis distances (BC), Unifrac distances (Uni), weighted Unifrac distances (wUni).
**A.** Comparison of soil-reared insects and control insects. Models contain one explanatory variable: soil treatment. **B.**
Comparison of soil-reared insects. Models contained two explanatory variables: site and soil subsample

**Figure 3:**
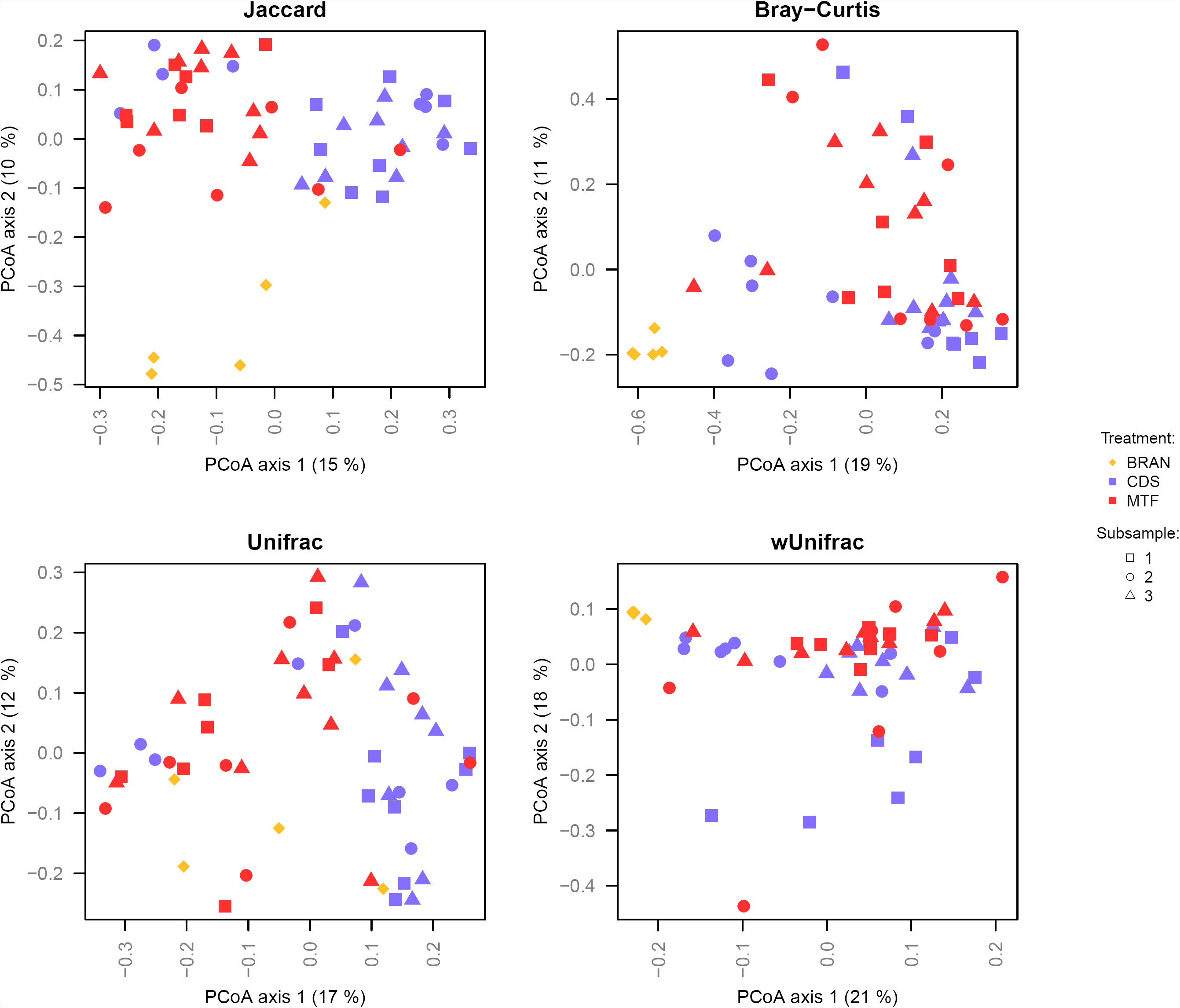
PCoA analysis based on the four beta diversity distances. Each dot corresponds to one insect. The percentage of variance explained by each axis is indicated in brackets. Yellow, blue and red dots correspond to BRAN (control), CDS and MTF samples respectively. For CDS and MTF samples, dot shape represents the identity of the soil subsample, i. e. CDS1, CDS2 and CDS3, or MTF1, MTF2 and MTF3.

The microbiota profiles of insects placed in soils from the same site (i.e. CDS or MTF) or in the same soil subsample (e.g. CDS1, CDS2 or CDS3) did not cluster together perfectly. However, a second PERMANOVA model for these samples identified two explanatory factors, soil sampling site (i.e. CDS or MTF) and subsample identity (e.g. CDS1, CDS2 or CDS3), as having a significant impact on gut community composition (Table 1B). Indeed, sample site explained 14 and 8% of the variance and soil subsample explained 17 and 18% of the variance, for the Jaccard and Bray-Curtis indices, respectively.

As reported above, the soil-reared insects had a microbiota dominated by *Enterobacteriaceae* (Figure 2C). We thus estimated Unifrac distances, which take into account the phylogenetic distances between OTUs, and wUnifrac distances, which also take relative OTU abundance into account. With these corrections, the differences between control insects and soil-reared insects were significant only when relative OTU abundance was taken into account (Figure 3; Table 1A). Subtle but significant effects of sample site and soil subsample on community composition were also observed with the Unifrac and wUnifrac indices (Figure 3; Table 1B).

Overall, our results show that soil treatment changes the community composition of the gut microbiota and that this change is detectable despite inter-individual variability. The bacterial communities present in the gut differ both between sample sites and between soil subsamples.

### Most of the changes in the microbiota concern the relative abundances of OTUs

We then pooled all individuals of a given treatment to determine which OTUs are found in at least one individual for each treatment. The 47 OTUs found in control insects were also present in the insects of the soil treatment groups (Figure 4A). The 44 OTUs common to all three conditions matched 97% of the reads for soil-reared insects (gray area in Figure 4B and Figure 4C). However, after soil treatment, *Pseudomonas*, the dominant OTU in control insects (98% of the reads) accounted for only 27 and 23% of the reads in CDS and MTF samples, respectively (Figure 4C). Conversely, *Serratia* species, together with the *Enterobacter* group, which accounted for less than 1% of sequence reads in controls, accounted for 35% and 43% of the reads for CDS and MTF, respectively.

**Figure 4:**
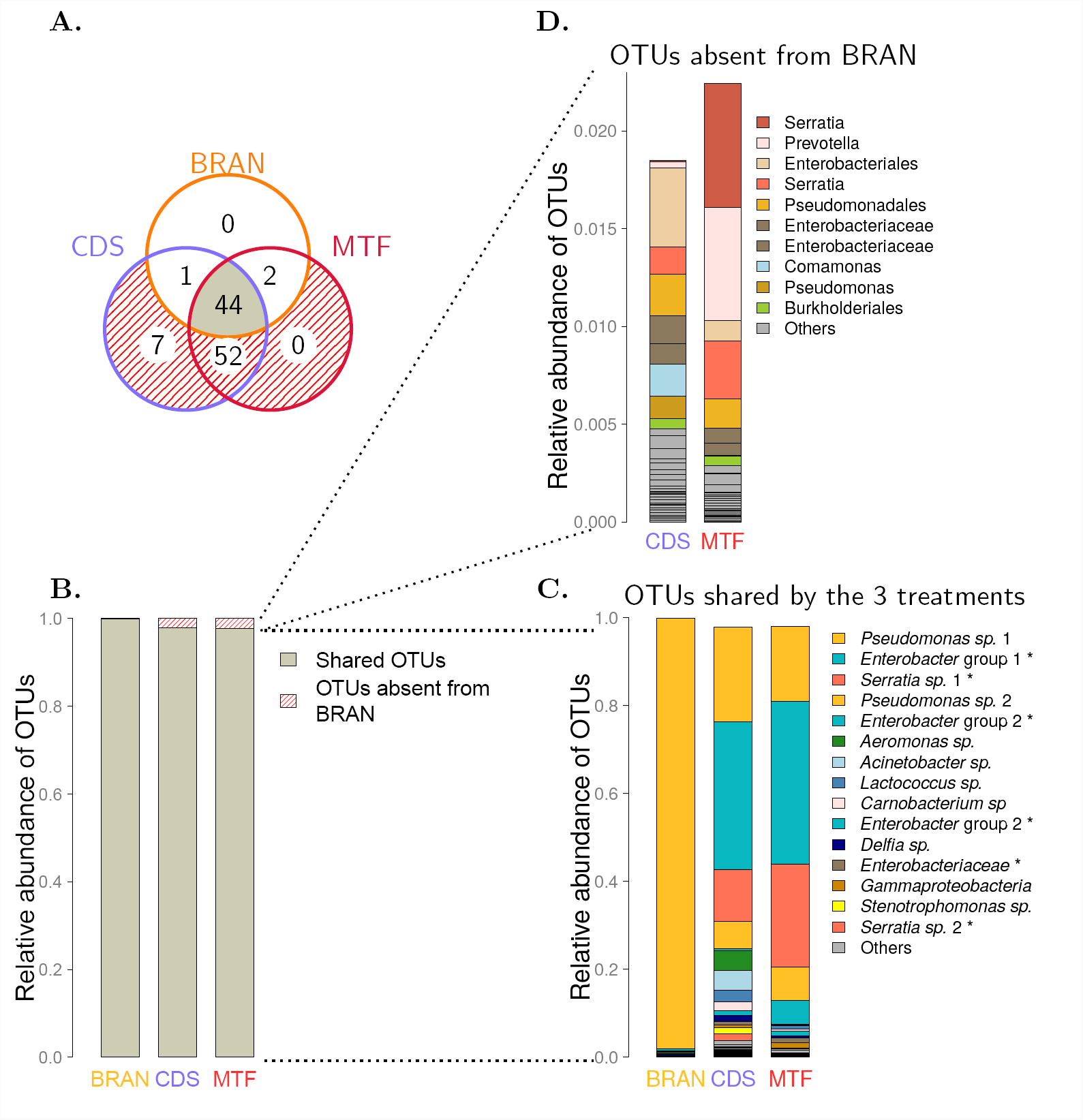
Assignment of shared OTUs according to the V3-V4 region of the 16S rRNA gene. **A.** Venn diagram of OTUs found in at least one insect from each treatment. **B.** Bar plot of the relative abundance of the 44 OTUs common to the three treatments (in gray) and the 59 OTUs found only in soil treatments (CDS and MTF) (red stripes). The taxonomic assignment of these OTUs is detailed in **C.** and **D.**. Insects from the various treatment were pooled for these bar plots: 5 insects for BRAN, 24 insects for CDS and 22 insects for MTF. The relative abundance of OTUs was calculated from the total number of reads for each insect pool. We show here taxonomic assignments to genus level or to the lowest taxonomic level, for which the bootstrap score was < 80%. Some OTUs differ in sequence, but were assigned to the same taxonomic group. These sequences are differentiated by a number. On each graph, the 15 OTUs with the largest relative abundance are shown in color and the others are grouped together in the “Others” category. OTU names followed by a star (*) belong to the *Enterobacteriaceae* family.

For confirmation of our initial metabarcoding results, we performed a second metabarcoding analysis with another marker, a 300 bp region of the *gyrB* housekeeping gene (see Additional File 1). This single-copy marker has been shown to provide assignments to more precise taxonomic levels than the 16S rRNA gene (Barret et al. 2015). In accordance with the results obtained with the 16S rRNA gene marker, *Pseudomonas* was the dominant OTU in control insects (more than 99 % of the reads) and its relative abundance was lower in soil-reared insects (CDS: 14 % MTF: 17 % of the reads). The genus *Serratia* and the *Enterobacter* group accounted for less than 0.06 % of the reads in control insects and a large proportion of those for the insects in the two soil treatment groups (CDS: 57 % MTF:70 % of the reads).

Finally, we also identified with 16S rRNA 59 OTUs that were not detectable in control insects but were present at low abundance (3% of the reads) in at least one soil-reared insect (red dashed area in Figure 4B and Figure 4D). These OTUs may correspond to taxa that were absent from the insects before soil treatment, and that colonized the insect gut during incubation in soil. Alternatively, they may have been present in the control insects at densities below the PCR detection threshold. Their abundance would then have increased above this threshold during incubation, just like the abundances of *Serratia* or *Enterobacter*. Overall, our data strongly suggest that the main effect of soil treatment is a change in the relative abundances of OTUs, although low levels of bacterial colonization from soil cannot be ruled out.

### The balance between members of the resident OTUs contributes to the variation of abundances after soil treatment

We assessed the variation of OTU balance after soil treatment further, by quantifying the bacterial taxonomic groups present in all treatments but with different relative abundances between the two contrasting sets of conditions studied (control versus soil-reared). We first characterized the gut resident microbiota in our larvae, as the OTUs present in at least 95% of our samples (following (Falony et al. 2016)). Based on 16S rRNA gene metabarcoding, we identified five resident OTUs: four *Enterobacteriaceae* (*Enterobacterericeae* 1, *Enterobacterericeae* 2, *Serratia* and *Enterobacter* group) and *Pseudomonas*. The resident OTUs obtained with the *gyrB* gene consisted of two OTUs, *Pseudomonas* and *Serratia*, confirming the existence of an invariant bacterial population in our insect gut microbiota. Based on the composition of this resident microbiota, we chose to monitor *Pseudomonas* and the *Enterobacteriaceae* to check for changes in the abundance of these bacteria following treatment. We performed quantitative PCR (qPCR) on a subset of 17 samples, including the five control insects and two insects for each soil subsample. We first calculated the gene copy number (GCN) of the 16S rRNA gene in each sample, using a universal primer pair targeting *Eubacteria* (uni16S primers). As the number of 16S rRNA gene copies varies across *Eubacteria* lineages (between 1 and 15 copies per genome, Lee et al. (2008)), the GCN cannot be used to quantify the number of bacterial cells with precision (Angly et al. 2014). However, in our samples, GCN/μL ranged from 10^7^ to 10^8^ and did not differ significantly between samples (Kruskal-Wallis rank sum test, chi squared = 2.66, df = 2, p-value = 0.26), which suggests that the total number of bacteria was similar in our 17 samples. We then targeted a region of the 16S rRNA gene specific to the *Pseudomonas* genus, (*Pse* -16S: 251 nucleotides of the V3-V4 hypervariable region, with 4 to 7 copies per genome Bodilis et al. 2012), and a region of the *rplP* gene, region specific to the *Enterobacteriaceae* family (Entero-*rplP* : 185 nucleotides of the *rplP* gene, one copy by genome). The *Pse* -16S GCN in soil-reared insects was one tenth that in control insects (Figure 5A). Conversely, the Entero-*rplP* GCN was 100 times higher in soil-reared insects (Figure 5B). Soil acclimation therefore seems to induce a decrease in *Pseudomonas* abundance and an increase in *Enterobacteriaceae* abundance. Our data suggest that the main effect of soil treatment is to modify the relative abundances of the resident bacterial communities of the gut microbiota.

**Figure 5:**
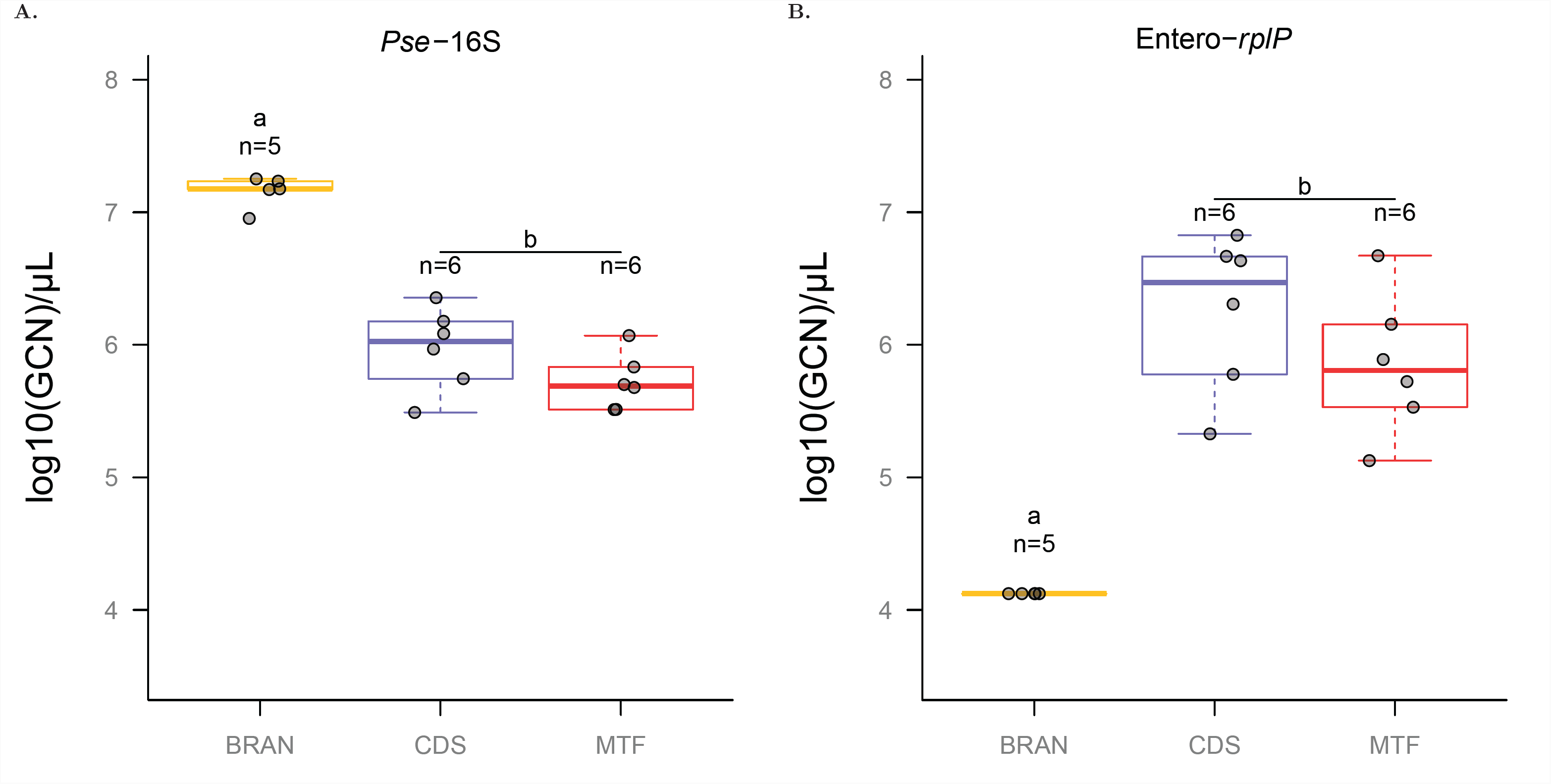
Quantitative PCR on two taxa of the core microbiota. **A.** Gene copy number (GCN) per μL of DNA extract for *Pse*-16S, a specific marker of the genus Pseudomonas. Pairwise Wilcoxon rank sum test with Holm p-value adjustment, BRAN-CDS: p-value = 0.013, BRAN-MTF: p-value = 0.013, MTF-CDS: p-value = 0.18. **B.** GCN per μL of DNA extract of Entero-*rplP*, a specific marker of the *Enterobacteriaceae* family. Samples from control (BRAN) had the maximum Ct value of 40, meaning that the initial Entero-*rplP* quantity was under the qPCR detection threshold, i.e. < 246 GCN. Pairwise Wilcoxon rank sum test with Holm p-value adjustment, BRAN-CDS: p-value = 0.016, BRAN-MTF: p-value = 0.016, MTF-CDS: p-value = 0.31.

## Discussion

Rearing larvae in soil rather than in bran caused major changes in gut microbiota structure. Soil-reared larvae have a richer and more diverse gut microbiota than control larvae. Despite considerable inter-individual variability, we found that the changes in community composition depended on both the site from which the soil was obtained, and the precise soil subsample used. An analysis of the OTUs found in the different samples suggested that the main effect of the soil treatment was a change in the relative abundance of OTUs. We confirmed this trend by qPCR for the two main taxonomic groups displaying changes in abundance: the *Enterobacteriaceae* family and the genus *Pseudomonas*, which predominated in soil-reared insects and in the control, respectively.

Our rearing conditions (laboratory versus soil acclimatization) were associated with two types of gut microbial patterns, consistent with previous findings for laboratory-reared and wild insects. On the one hand, gut microbiota communities of laboratory-reared insects, which are usually maintained on very simple media and diets, are dominated by one or two bacterial strains: *Pseudomonas* in our study, *Enterococcus* in moths (Chen et al. 2016; Staudacher et al. 2016) or the *Enterobacteriaceae* group *Orbus* in fruit flies (Chandler et al. 2011). On the other hand, following soil treatment, our larvae harbored more complex community profiles, with several *Enterobacteriaceae* together with the *Pseudomonas* strain that we found in control insects. Wild coleopterans, such as the forest cockchafer, *Melolontha hippocastani*, which has a soil-dwelling larval stage, have a microbiota dominated by *Enterobacteriaceae*, essentially a consortium of *Serratia*, and a Shannon diversity index close to that observed here for soil-reared insects (Arias-Cordero et al. 2012). Other coleopterans, such as *Agrilus planipennis* and *Nicrophorus vespiloides* (Vasanthakumar et al. 2008; Wang and Rozen 2017), both sampled from the wild and reared on a natural diet, also have microbiotas dominated by *Pseudomonas sp.*, the *Enterobacter* group and *Serratia sp.*. These findings suggest that our protocol can be used to mimic soil-dwelling insects effectively with reared insects. This might make it possible to obtain large numbers of individuals while working on a relevant set of bacteria in further studies of the insect gut microbiota. Moreover, we focused here on the gut microbiota, but soil treatment probably modifies the entire microbiota, including the cuticular bacterial community. Our methodology is therefore likely to be of particular interest for holobiont studies (Bordenstein and Theis 2015) involving controlled hypothesis-driven experiments on insects with a relevant total bacterial community.

The changes we observed in gut microbiota structure may result from major changes in insect diet, as insects may have access to different sources of food when incubated in soil compared to sterile bran. Our results fit well to the diet influences on microbiota documented in several *Drosophila* species (Chandler et al. 2011; Staubach et al. 2013; Vacchini et al. 2017), omnivorous cockroaches (Pérez-Cobas et al. 2015), termites (Mikaelyan et al. 2015), lepidopterans (Broderick et al. 2004; Belda et al. 2011; Priya et al. 2012) and a few coleopterans (Colman et al. 2012; Jung et al. 2014; Franzini et al. 2016; Kim et al. 2017). Changes in microbiota structure could also depend on physiochemical properties of the insect gut. In wood-feeding cockroaches, different parts of intestinal tract showed differences in pH, redox potential and hydrogen contents, and were associated to different bacterial communities (Bauer et al. 2015). The ingestion of soil particles probably modifies some of these properties of the gut. The fact that the soil characteristics differed between the two sampling sites (low sand/silt ratio for Causse-De-La-Selle (CDS), and higher sand/silt ratio for Montferrier (MTF)) could thus explain in part their different impacts on *T. molitor* gut microbiota.

The changes in the gut bacterial population may depend not only on treatment, but also on the bacterial community initially present in the gut. Previous studies (Jung et al. 2014; Osimani et al. 2018) showed that a *Spiroplasma* species predominated in the gut microbiota of the larval lineage, even after and environmental change. *Spiroplasma* has been shown to be a heritable endosymbiont in *Drosophila* (Mateos et al. 2006). Similar effects were observed for other endosymbionts, such as *Wolbachia, Cardinium, Blattabacterium*-like and putative *Bartonella*-like symbionts in mites *Tyrophagus putrescentiae* following dietary changes (Erban et al. 2017). In all these case, endosymbiont seem to impede major shifts in the gut microbiota or conceal changes in frequencies that may occur in low-abundance OTUs. This effect is absent in our experiment, probably because the insects we used are associated to *Spiroplasma* or any other endosymbiotic bacteria.

Our results also provide interesting insight into the spatial variation of the gut bacterial community in insect populations. The differences observed after incubation in soil from different plots were consistent with the findings of other studies on coleopterans, in which the dissimilarity of the gut bacterial community between individuals is correlated with the distance between sampling sites (Adams et al. 2010). However, we also observed a difference in the gut microbiota between insects incubated with soils collected a few meters apart, at the same sampling site, and this difference was detectable despite high levels of inter-individual variation. Minor environmental differences thus have a detectable impact on the gut microbiota and structure this microbiota within insect populations over very small geographic scales.

Overall, our experiments indicate that gut microbiota can be readily changed by modifying the environment in which the insects are living. We identified resident taxa present in all the environments we tested. These taxa change in relative abundance with environmental changes. The range of environmental conditions tested here is narrower than that experienced by insects in the wild, but results suggest that, following changes in environmental conditions, the insect gut microbiota maintains a stable composition, but displays plasticity in terms of its structure.

## Availability of data and material

Both the 16S rRNA and *gyrB* datasets generated and analyzed in this study are available from the ENA (European Nucleotide Archive) repository, http://www.ebi.ac.uk/ena/data/view/PRJEB21797

## Funding

MC obtained PhD funding from the French Ministry of Higher Education, Research and Innovation. Metabarcoding sequencing was funded by the MEM-INRA metaprogram (P10016). This work was also supported by the French Laboratory of Excellence project “TULIP” (ANR-10-LABX-41; ANR-11-IDEX-0002-02)

## Authors’ contributions

MC, JBF and SG conceived the study. MC designed and performed the experiments. AL performed qPCR analysis. MC and JCO analyzed the data. JBF and SG supervised the project. All authors wrote, read and approved the final manuscript.

## Acknowledgments

We thank Marie Frayssinet for help with soil sampling and insect acclimatization, and Lucie Zinger for help with data analysis.

## Additional Files

### Additional file 1: Relative abundance and taxonomic assignment of OTUs according to the *gyrB* gene

Insects from the various treatments were pooled for these bar plots: 5 insects for BRAN, 24 insects for CDS and 22 insects for MTF. The relative abundance of OTUs was calculated from the total number of reads for each insect pool. We show here taxonomic assignments to genus level or to the lowest taxonomic level for which the bootstrap score was > 80%. Some OTUs differ in sequence but were assigned to the same taxonomic group. These sequences are differentiated by a number. On each graph, the 15 OTUs with the largest relative abundances are shown in color and the others are grouped together in the “Others” category. OTU names followed by a star (*) belong to the *Enterobacteriaceae* family.

### Additional file 2: Example of a microbiota pattern in PCR replicates

We checked the reproducibility of PCR, by performing three technical PCR replicates (the three bars of the chart) on a sample chosen at random, with the whole metabarcoding procedure performed separately for each replicate. We show here the results for the CDS1D3 sample.

### Additional file 3: OBITools workflow for 16S rRNA analysis

RMD_OBITools_workflow_V3V4.pdf and RMD_OBITools_workflow_gyrB.pdf contain OBITools, bash and R scripts used to obtain the OTU abundance table from raw sequencing data for both the 16S rRNA and *gyrB* genes.

### Additional file 4: Raw table of reads counts

tab_div_V3V4.csv and tab_div_gyrB.csv contain raw abundance data and diversity indices for each sample, as determined with the 16S rRNA and *gyrB* genes, respectively. Samples are shown in rows and OTUs in columns.

### Additional file 5: OTU taxonomic assignment

V3V4_assignment.txt is the assignment data for each 16S rRNA OTU obtained with RDPclassifier and the RDPII database. gyrB_assignment.csv is the assignment data for each *gyrB* OTU obtained with the MOTHUR classifier and the Barret et. al 2014 reference database.

### Additional file 6: Primers used for qPCR

PE_*standard*_ corresponds to PCR efficiency on gDNA standard samples, PE_*gut*_ corresponds to PCR efficiency on a pool of gut DNA from samples used for qPCR analysis.

### Additional file 7: Statistical analysis workflow

RMD_R_workflow.pdf contains R scripts used to perform statistical analysis and to produce the figures presented in this study.

### Additional file 8: R functions used in the statistical analysis workflow

- src_routine_boostrap_threshold. R is an R function for extracting the lowest taxonomic level according to a given bootstrap threshold from assignment files

